# Attention and working memory investigated through the P300 component in children practicing Karate at different stages of biological maturation

**DOI:** 10.1101/2024.08.18.608511

**Authors:** Eduardo Nicoliche, Alexandre Vasconcelos, Marcos Machado, Guaraci Tanaka, Renan Vicente, Adriana Marques, Élida Costa, Mariana Gongora, Jessé Di Giacomo, Marcelo Nobre, Silmar Teixeira, Victor Hugo Bastos, Mauricio Cagy, Isabelle Fernandes, Caroline Machado, Pedro Ribeiro, Daya S. Gupta, Bruna Velasques, Henning Budde

## Abstract

**Aim:** To investigate attention and working memory, comparing children practice Karate and non-Karate practitioners at different stages of biological maturation through the amplitude and latency of the P300 component during the execution of a Go/No-Go paradigm.

**Material and Methods:** The P300 was analyzed for Fz, Cz, and Pz electrodes in 80 participants separated in two groups: an Karate practitioners group comprising Karate practitioners and comprising non-Karate practitioners. Each group was further subdivided according to the biological maturation range defined by Peak Height Velocity. In addition, the participants performed a Go/No-Go paradigm to measure amplitude and latency.

**Results:** The EEG analysis showed Ffr electrodes Pz and Cz, an interaction was found between group and Peak Height Velocity for the amplitude variable (respectively: F = 45.858; d = 0.38; p < 0.001 / F = 10.411; d = 0.17; p = 0.004). For the Fz electrode, a main effect was found between group and Peak Height Velocity (respectively: F = 40.330; d = 0.34; p = 0.010 / F = 36.730; d = 0.30; p = 0.012) for the variable amplitude and latency. main effect between group and Peak Height Velocity (respectively: F = 7.719; d = 0.14; p = 0.012 / F = 38.370; d = 0.31; p = 0.010).

**Conclusions:** In general, it is possible to conclude that participants in the Karate practitioners group exhibited electrocortical measures corresponding to greater efficiency in decision-making and attention processes, motor planning, working memory, attention allocation, motor execution, and greater attentional engagement. It was also demonstrated that, despite the children being at very close chronological ages, their biological maturation differed.

## 1. Introduction

The existing literature includes numerous studies that suggest an association between exercise and the development of cognitive functions in children and adolescents (1–3). Research on the impact of a single exercise session on cognitive functions suggests positive effects on inhibitory control, working memory, and cognitive flexibility (4–6). These effects appear to be modulated by exercise intensity, duration, time, and stage of development (5,7,8). A recent review of longitudinal studies indicates that individuals who engage in chronic exercise intervention demonstrate improved cognitive performance (9).

The biological maturation process encompasses a range of and neurophysiological changes that occur at varying rates for each individual, resulting in differences in motor performance (10). Biological maturation is a process of physiological and neurophysiological changes that occur at different speeds for everyone, resulting in differences in motor performance (11,12). In the context of exercise a classification based on maturation is necessary, as highlighted by research on children and adolescent (11,13,14). As a specific region of the brain matures, the individual displays behaviors corresponding to that area, provided that the relevant function is stimulated (15–17). Therefore, the phase of biological maturation serves as a critical indicator of the relationship between cognitive and motor skills in children who engage in sports activities (18).

Quantitative electroencephalography (EEGq) is a non-invasive and method with high temporal resolution that investigates cortical regions involved in attention and working memory processes, quantifying electrical activity through analysis in both the time and frequency domains (19). Event-Related Potentials (ERP) can explore distinct aspects of stimulus processing in Go/No-Go paradigms and serve as a technique for observing the mechanisms involved in cognitive functions, providing information about the processes between observation of a stimulus and execution of a response. One of these is the positive wave called P300, which is evident between 300 and 600 ms after viewing the stimulus (20,21). This wave reflects various processes related to the allocation of attention to the presented stimulus (22–24).

Little research has investigated the effect of physical exercise, especially Karate, on children’s attention and working memory using the classification of maturational stages and neuroimaging equipment. For this reason, the literature presents conflicting results, since individuals may be within the same age range, but at different maturational stages, making performance and cognitive capacity different. Therefore, The aim of this research was investigate attention and working memory, comparing children practice Karate and non-Karate practitioners at different stages of biological maturation through the amplitude and latency of the P300 component during the execution of a Go/No-Go paradigm. The study hypothesizes that lower P300 amplitude and latency will be observed in children who practice Karate and are at a more advanced maturational stage, because martial art promotes the improvement of specific movements related to motor control, balance, coordination, agility and the complexity of movements combined without obeying an organized flow of movement, events are random and movements need to adapt to new contexts.

## 2. Material and methods

### 2.1. Study design

This is a cross sectional comparison research.

### 2.2. Participants

The sample consisted of 80 participants, divided into two groups: comprising Karate practitioners (n = 40) and a comprising non-Karate practitioners (n = 40). Each group was further subdivided according to the biological maturation range defined by Peak Height Velocity (28). Each group was subdivision related to the biological maturation range according to Peak Height Velocity (28): Level −3 (n = 20) and Level −2 (n = 20) (figure 1). Inclusion criteria encompassed participants aged between 9 and 17 years, with right-handed laterality, who were enrolled in physical education classes at school. Individuals with any neurological or psychological disorder were excluded from the study. Additionally, those who engaged in systematic training that is not karate more than once a week were also excluded

**Figure 1.**
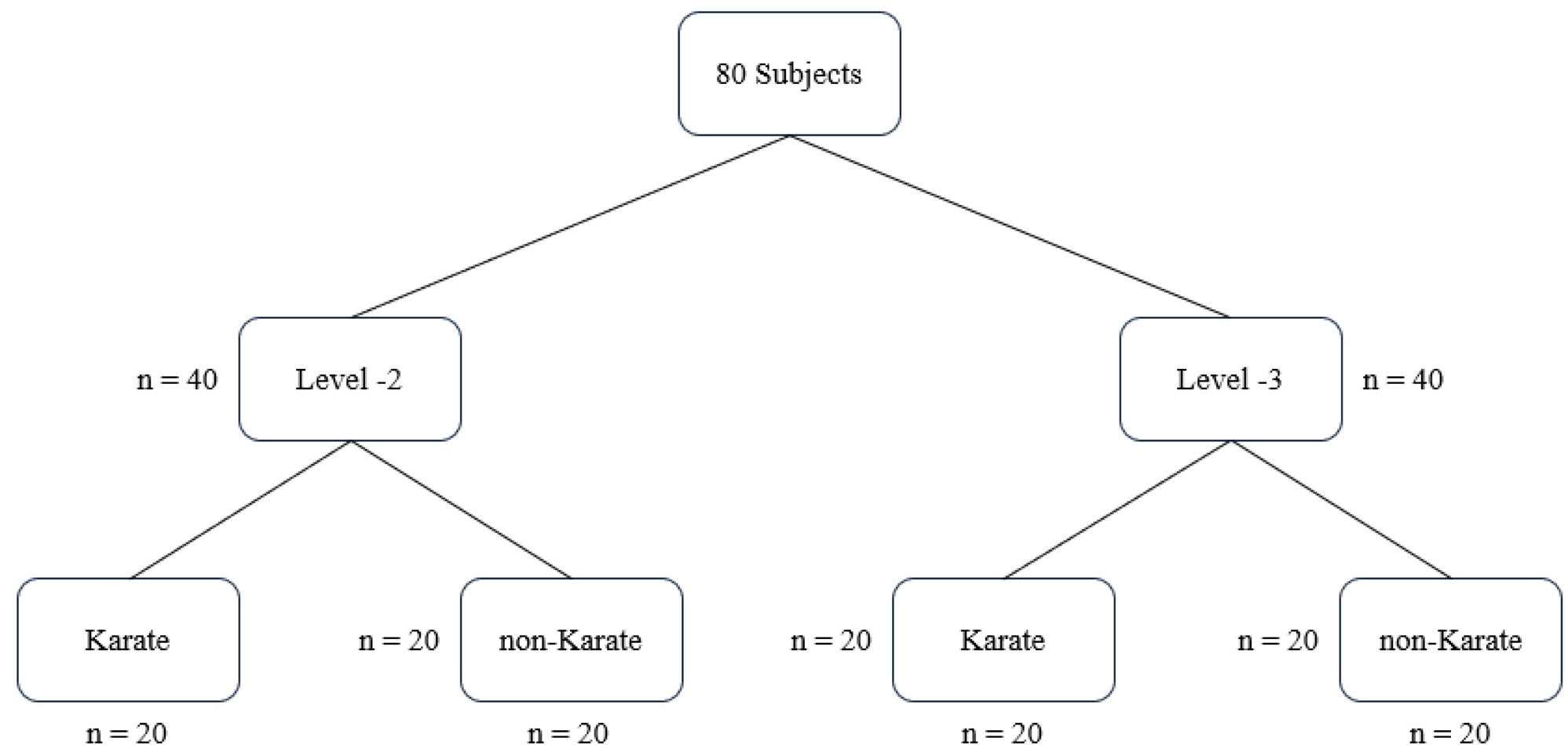
Division of groups

Furthermore, the Edinburgh Handedness Inventory was applied to verify the predominance of the right hand (25).

All participants and their adult guardians signed a Free and Informed Assent Form and a Free and Informed Consent Form. These forms detailed the experiment’s objectives and the experimental conditions to which the subjects would be exposed.

The research was conducted in the Laboratory of Neurophysiology and Neuropsychology of Attention (LANNA), following the Declaration of Helsinki and approved by the local Ethics Committee.

### 2.3. Peak Height Velocity (PHV)

Biological maturation is related to the child’s brain development and it is important to define whether the attentional process is in accordance with the maturation process (17,26,27). There are different methods for evaluating biological maturation (28). Peak Height Velocity is based on measurements of anthropometric body proportions (29), being one of the most used measurement methods in studies investigating biological maturation (11,29,30). Each participant’s PHV was estimated using height, sitting height and body mass using the methods of Mirwald et al. (29). The classification was carried out according to Machado et al. (16).

### 2.4. Karate Training

All participants in the Karate group had been training for 6 months, two sessions per week, lasting approximately 60 minutes per session. The training was always conducted by the same instructor. The group of non-karate practitioners did not practice other sports and did not differ in BMI from the group of practitioners.

### 2.5. Experimental procedure

The participants were individually referred to the laboratory, where they underwent anamnesis, anthropometric assessment to calculate Peak High Velocity and the experimental paradigm Kanizsa Figures with electroencephalographic capture.

The experimental paradigm was conducted in an environment with reduced lighting and acoustic insulation to minimize sensory interference. The subjects were seated in a comfortable chair with their right arm rested on support to minimize muscular artifacts. Subsequently, the Kanizsa Figures experimental protocol was carried out according to Sokhadze et al. (31), which consisted of three minutes of rest, followed by five stages of performing the paradigm, (five minutes each stage) and, finally, another three minutes of rest.

### 2.6. Paradigm Figures of Kanizsa

The paradigm comprises a series of stimuli presented randomly, with one presented infrequently. The subjects were instructed to discriminate the target stimulus (“Go” - infrequent) from the non-target stimuli (“No-Go” - frequent). The stimuli are the Kanizsa square (target), the Kanizsa triangle, the non-Kanizsa triangle, and the non-Kanizsa square (Figure 2). The non-target Kanizsa triangle is introduced to differentiate the processing of Kanizsa figures and targets, figures of the stimuli available in Sokhadze et al. (31). The stimuli comprise three or four inducing discs, that are considered the shape feature, and they constitute an illusory figure (square or triangular) or not (characteristic of collinearity) (31). The participants were instructed to maintain constant attention on the screen and to respond as quickly as possible when the target figure appeared.

**Figure 2.**
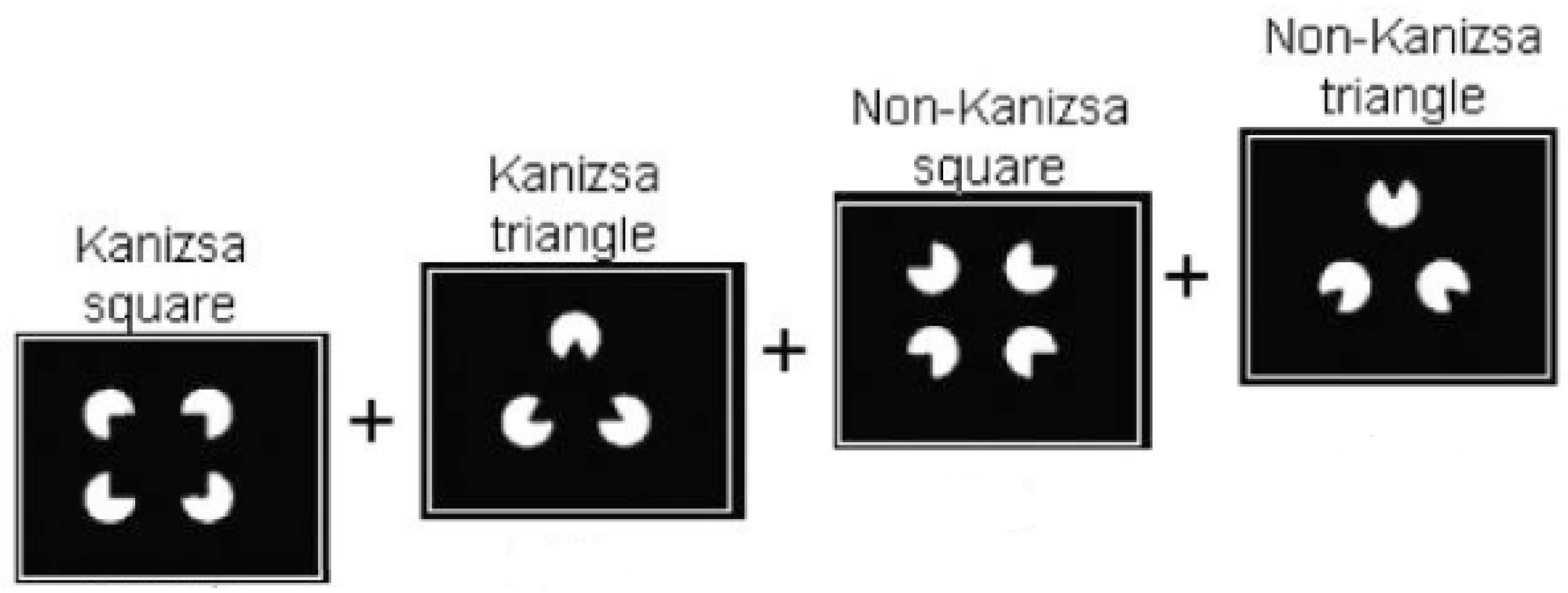
Kanizsa figures

Sokhadze et al. (31)

### 2.7. Behavioral parameter – Reaction Time (RT)

The behavioral parameters were extracted from the Reaction Time (RT) discriminated from the target stimulus of the Kanizsa Figures paradigm (31).

### 2.8. EEG recording and data processing

To capture the electroencephalographic signal, the BNT-36 device (EMSA – Equipamentos Médicos, Brazil) with a system that uses an analog-to-digital (A/D) converter board with 16-bit resolution, in conjunction with the ERP_Aquisition acquisition software, responsible for both signal acquisition and the generation and presentation of stimuli on the computer screen. Its configuration uses 60 Hz Notch digital filtering, 0.1 Hz high-pass, and 100 Hz low-pass filters, with 400 samples per second sampling rate.

The international 10-20 system was used to place 20 monopolar electrodes along the scalp and one electrode on the lobe of each ear (average auricular reference) (32,33). These electrodes were pre-attached to a nylon cap (ElectroCap Inc., Fairfax, VA) and used according to each subject’s cranial perimeter. The impedance levels of each electrode were kept below 5 kΩ.

The ERP waveforms were estimated using a coherent average from the electroencephalographic data collected during the experiment and processed using routines developed in LANNA. Each epoch used for both analyses consisted of EEG excerpts beginning 1,000 ms before the onset of the visual stimulus and ending 1,000 ms after the onset of the stimulus. The analysis excluded any data that exhibited muscle artifacts or gross eye movements. The ERPs were classified according to the nature of the target stimuli (“Go”).

### 2.9. Statistical analysis

The factors Group (Karate × Control) and Peak Growth Velocity (Level-2 × Level −3) were analyzed.

The analysis was performed by comparing the time domain parameters (ERP) of the P300 component: Amplitude and Latency.

First, the raw data was plotted to identify its variability and extract outliers. The Kolmogorov-Smirnov test was used to evaluate the normality of the distribution of the investigated parameters (P300 Amplitude and Latency) (p = 0.65). Then, a “two-way” ANOVA (Group × PHV) was conducted for the Fz, Cz, and Pz electrodes to separately examine the cognitive and electrophysiological aspects of each cortical region. Bonferroni correction adjusted probability (p) values due to the increased risk of a type I error when doing multiple statistical tests. A t-test was applied to perform inspect interactions when necessary. The effect size analysis was realized by the Cohen’s d. The effect sizes were calculated (≤0.039: no effect, 0.04–0.24: minimum, 0.25–0.63: moderate, ≥0.64: strong according to Ferguson (34). The same parameters were maintained for the analysis of Reaction Time. The significance adopted was p ≤ 0.05. All analyses were conducted using SPSS for Windows version 20.0 (SPSS Inc., Chicago, IL, USA).

## 3. Results

### 3.1. Reaction time

An interaction was found between the factors Group and PHV (F = 65.644; d = 0.55; p < 0,001). Interaction inspection found a significant difference in PHV (p < 0.001) (figure 3). The reaction time values are shown in table 1.

**Figure 3.**
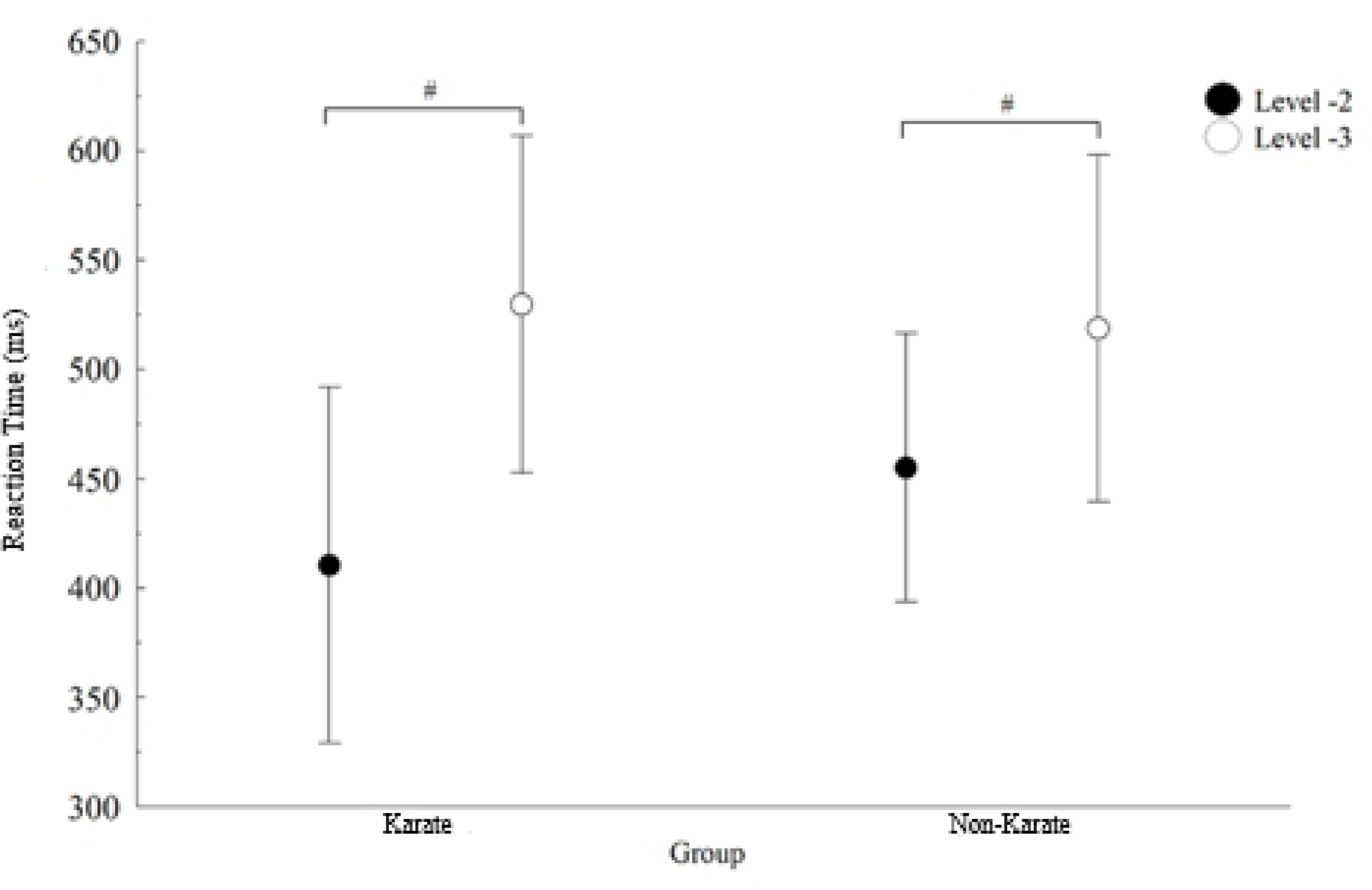
Inspection of interaction between Group and PHV for reaction time. #: significant difference for PHV.

**Table 1.** Reaction time values.

| *Reaction time (ms)* | *Karate practitioners* |  | *non-Karate practitioners* |  |
| --- | --- | --- | --- | --- |
|  | **PHV Level -2** | **PHV Level -3** | **PHV Level -2** | **PHV Level -3** |
| | 412.8 ± 73.25 | 528.3 ± 72.88 | 459.1 ± 61.73 | 525.6 ± 75.40 |

### 3.2. Latency and amplitude of P300 components

The amplitude and latency values of the P300 component arer shown in table 2. The latency and amplitude of the P300 components were analyzed using “two-way” ANOVA to compare the factors Group × PHV at electrodes Fz, Cz, and Pz. The ERP graphs for Fz, Cz, and Pz are presented in Figure 4.

**Table 2.** Amplitude and latency values of the P300 component.

| Electrode | *Karate practitioners* |  |  |  | *non-Karate practitioners* |  |  |  |
| --- | --- | --- | --- | --- | --- | --- | --- | --- |
|  | PHV Level -2 |  | PHV Level -3 |  | PHV Level -2 |  | PHV Level -3 |  |
| | Amplitude ($\mu V$) | Latency (ms) | Amplitude ($\mu V$) | Latency (ms) | Amplitude ($\mu V$) | Latency (ms) | Amplitude ($\mu V$) | Latency (ms) |
| Pz | $7.0 \pm 1.80$ | $532.5 \pm 52.63$ | $11.2 \pm 2.12$ | $551.2 \pm 51.43$ | $13.3 \pm 2.82$ | $528.6 \pm 57.42$ | $14.8 \pm 2.20$ | $546.0 \pm 55.87$ |
| Cz | $6.7 \pm 1.82$ | $532.1 \pm 56.22$ | $8.1 \pm 1.77$ | $573.3 \pm 52.34$ | $8.6 \pm 2.13$ | $548.6 \pm 56.4$ | $9.8 \pm 2.37$ | $583.3 \pm 57.76$ |
| Fz | $4.8 \pm 1.33$ | $623.4 \pm 58.72$ | $2.2 \pm 0.71$ | $553.1 \pm 55.73$ | $3.7 \pm 1.10$ | $642.3 \pm 62.37$ | $8.9 \pm 2.11$ | $584.6 \pm 59.22$ |

**Figure 4.**
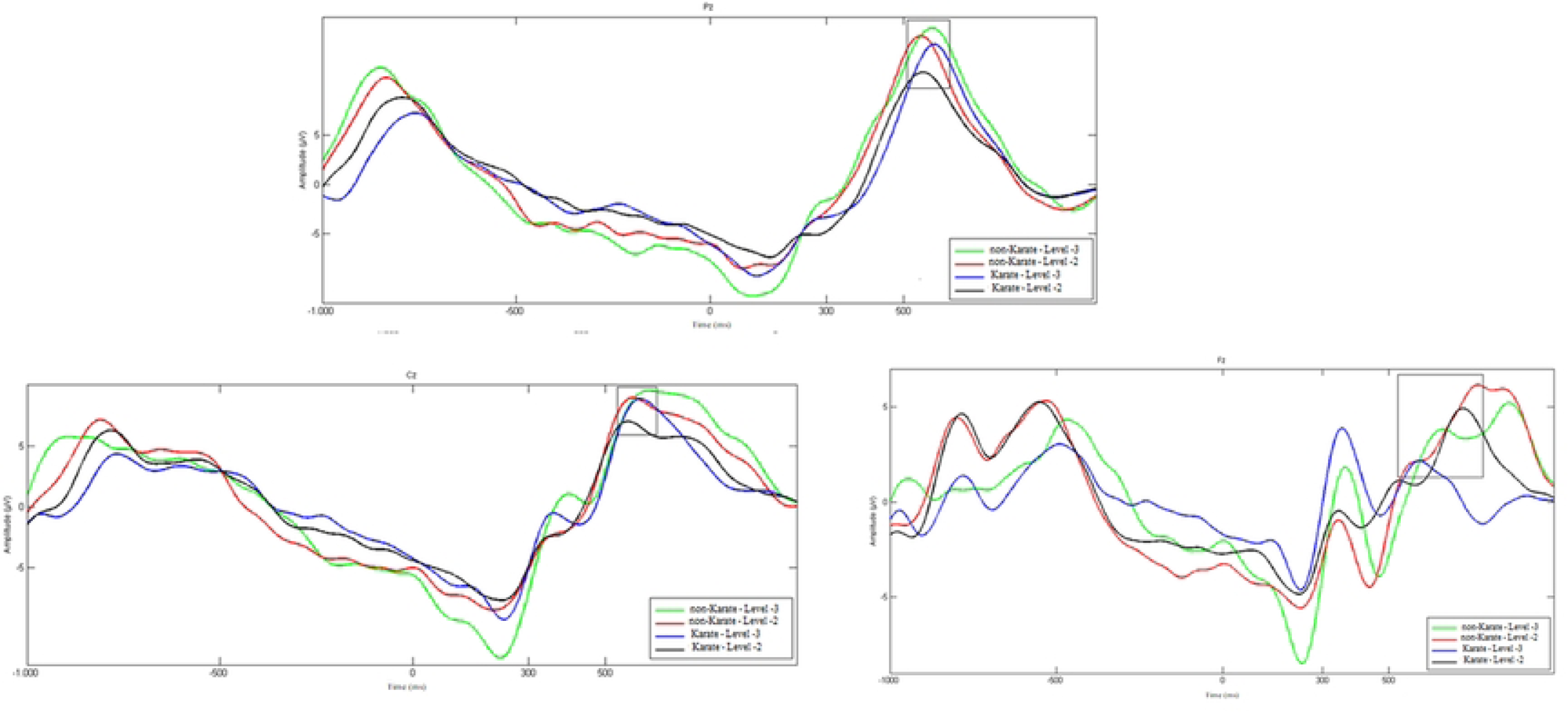
Representation of the Event-Related Potential for the Pz, CZ and Fz electrode

For the P300 amplitude variable, at the Pz electrode, an interaction was found between the factors Group and PHV (F = 45.858; d = 0.38; p < 0.001). Interaction inspection found a significant difference between groups (p = 0.005) and between PHV (p = 0.040) (Figure 5A). At electrode Cz, an interaction was found between the factors Group and PHV (F = 10.411; d = 0.17; p = 0.004). Interaction inspection found a significant difference in PHV (p = 0.006) (Figure 5B). At the Fz electrode, a main effect was found for the factors Group (F = 40.330; d = 0.34; p = 0.010) (Figure 5C) and PHV (F = 36.730; d = 0.30; p = 0.012) (Figure 5D).

**Figure 5.**
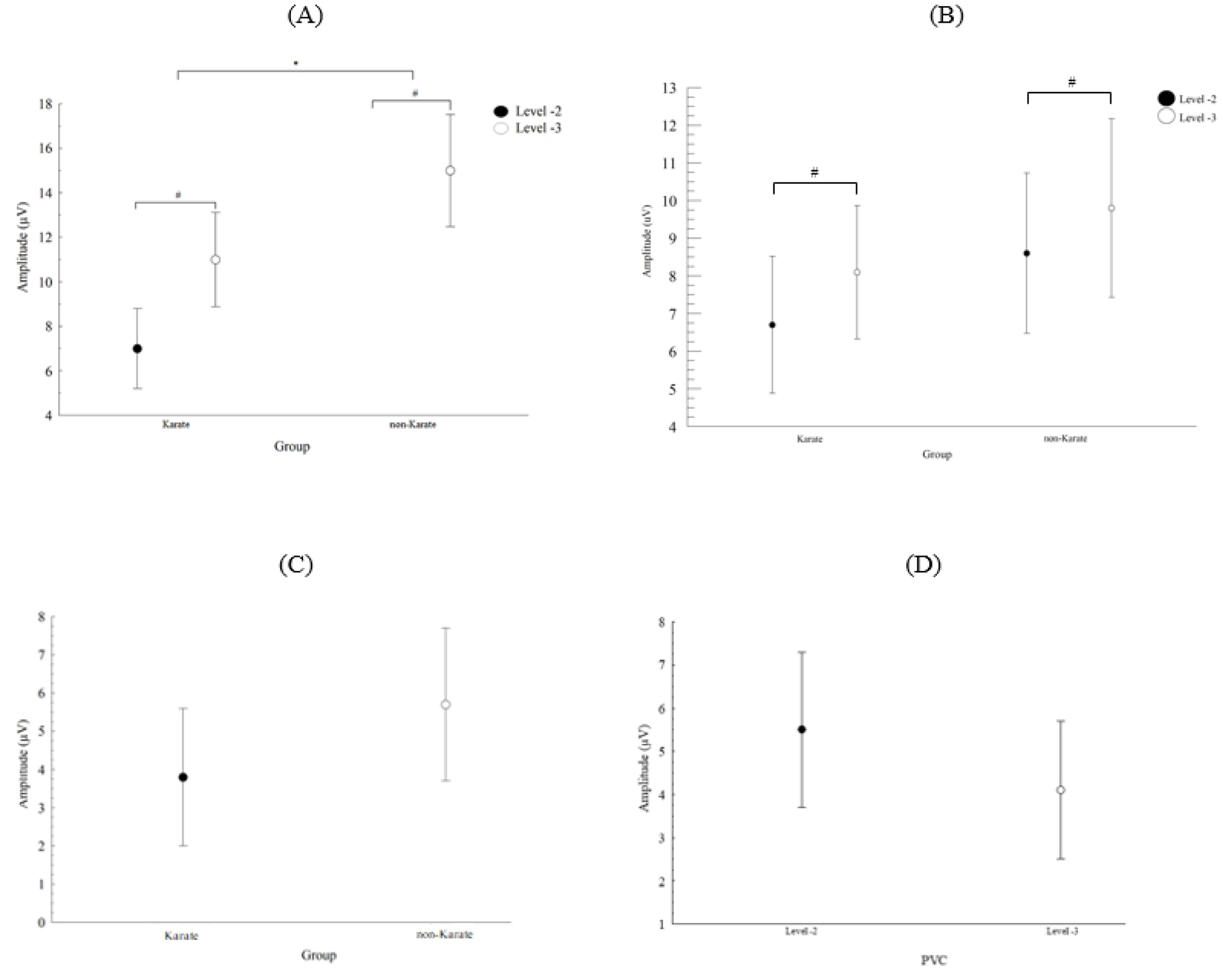
Inspection of interaction between Group and PHV at electrode Pz (A), nspection of interaction between PHV at electrode Cz (B) and Main effect for Group (C) and PHV (D) at electrode Fz on the P300 amplitude variable *: significant difference for Group. #: significant difference for PHV.

For the P300 latency variable, at the Fz electrode, a main effect was found for the Group factor (F = 7.719; d = 0.14; p = 0.012) (Figure 6A) and the PHV factor (F = 38.370; d = 0.31; p = 0.010) (Figure 6B). At electrodes Pz and Cz, no main effects or interactions between factors were found.

**Figure 6.**
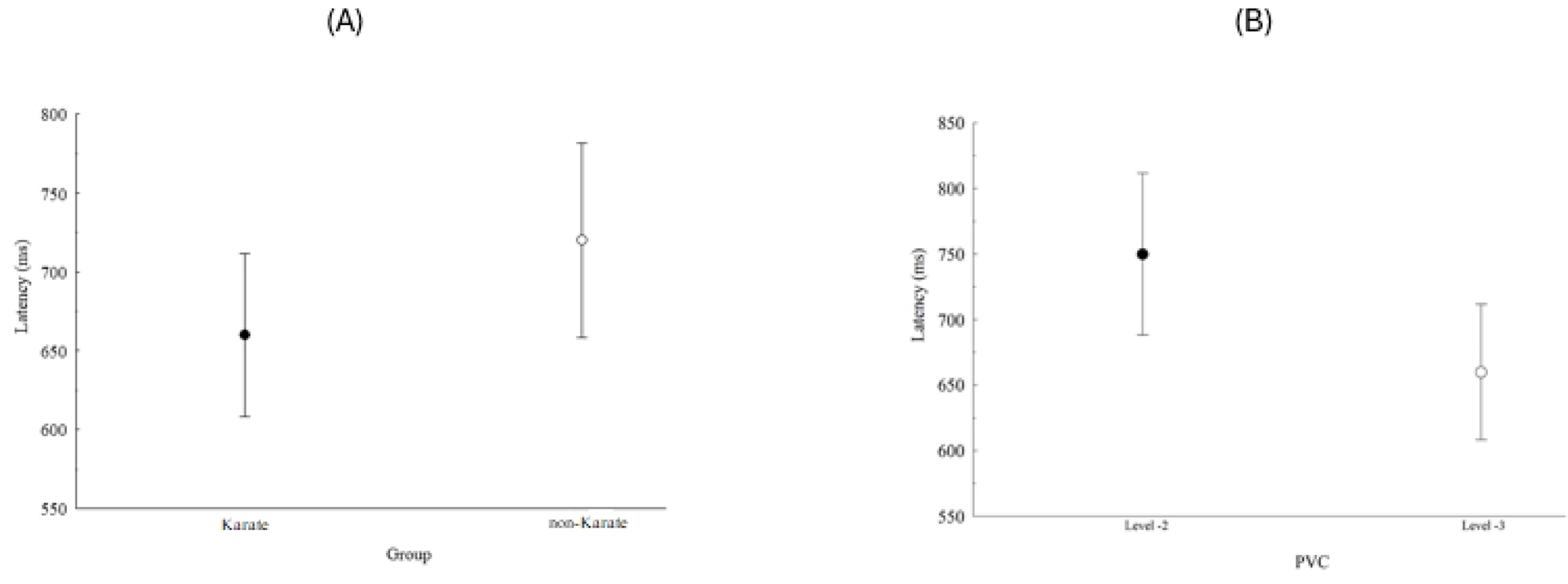
Main effect for Group (A) and PHV (B) on the P300 latency variable, at electrode Fz

The black rectangles represent the location, in relation to the time window, that was used to calculate the P300 latency and amplitude.

## 4. Discussion

The present study aimed to investigate attention and working memory, comparing children practice Karate and non-Karate practitioners at different stages of biological maturation through the amplitude and latency of the P300 component.

To achieve this, changes in the latency and amplitude of the P300 component were investigated during the execution of the Kanizsa Figures paradigm. Therefore, the discussion of the electrophysiological results for P300 amplitude and latency was divided into Frontal Cortex and Parietal Cortex: cognitive and sensorimotor integration (Pz); Central Cortex: motor preparation and execution (Cz); Executive Functions (Fz).

### 4.1. Behavioral Perspective – Reaction Time

The results of the present study demonstrated superior task performance, as measured by reaction time (RT), in participants with more advanced biological maturation (PHV Level −2). Additionally, the Karate practitioners group exhibited slightly superior performance, although this difference was not statistically significanthow is the effect size? compared to the non-Karate practitioners. Mezzacapra (35) used the “flanker task” to evaluate the RT of children from 6 years of age annually until they turned 9. The authors found lower RT as the years went by. Other research also used the “flanker task” with children of different ages, finding lower RT values in the decreasing order of age of the groups (36,37). In a subsequent study, Reuter et al. (10) carried out a study with participants aged between 20 and 81 years. The results indicated that children showed a slower RT than young and middle-aged participants; in the elderly, the RT was lower than in all age groups. Rueda et al. (38) compared children between 8 and 12 years old with adults between 30 and 40 years old. The results indicated that adults exhibited shorter reaction times. These studies demonstrated that reaction time improves with age, thus supporting the hypothesis that humans improve the speed and efficiency of processing and cognitive control throughout their development into adulthood, with the decline observed only in adults older. These data are consistent with our findings. However, a significant discrepancy was observed between the two PHV classifications (Level −2 and Level 3).

Our data revealed no discernible difference between the non-Karate practitioners and Karate practitioners groups. In a study by Lima et al. (39), children who practiced Karate and non-practitioners between 8 and 11 years old were compared. RT was measured using a visuomotor paradigm, where participants had the numeral “1” as the target stimulus, and the distracting stimuli were the numerals “2”, “3,” and “4.” A lower RT was found in the group of Karate practitioners. Sharma et al. (40) corroborates the previous study by comparing the RT measured by a Go/No-Go sound stimulus paradigm between practitioners of different sports and non-practitioners aged between 10 and 19 years, finding lower RT among practitioners. Ludyga et al. (41) found similar results like the two previous studies when evaluating RT measured using a Go/No-Go characteristic visuomotor paradigm in children between 9 and 13 years old with at least six months of Judo practice with non-practitioners. Possibly, the Karate practice time needed to be longer for significant differences to be observed between the Karate practitioners and non-Karate practitioners groups. The heterogeneity of the experimental methods used in the reviewed research makes it difficult to compare results and better understand the topic directly.

### 4.2. Electrophysiological perspective

The data from this study were analyzed in the time domain using the Event-Related Potential (ERP) technique. The P300 component was selected due to its association with the final stage of information processing, reflecting working memory, attention, and the speed of information processing involved in the decision-making process (42). Changes in P300 amplitude and latency are correlated with mental processes to deal with a sequence of repetitive stimuli interrupted by a target stimulus (23,43). Therefore, the electrophysiological results for P300 amplitude and latency were discussed based on the interaction between the independent factors and main effects: Group and PHV.

### 4.3. Associative Somatosensory Cortex: Brodmann Area 7 (Pz)

At the Pz electrode, we observed interaction in the P300 amplitude between the factors: Group and PHV. Previous studies have identified that the P300 amplitude value recorded in this region indicates the ability to allocate attention and working memory resources, while latency reflects the ability to detect the target stimulus (44–46). Polich (23) and Lind et al. (47) statedthat P300 amplitudes are highest over the central parietal site. Thus, changes observed in the P300 amplitude in the Karate practitioners group, particularly in those closest to the PHV (Level −2), suggest an improvement in information processing for detecting and responding to target stimuli, these changes indicate better attention allocation and working memory, attributable to both the practice of Karate and their more advanced stage of development.

Lind et al. (47) identified higher latency values in children who play football and found no significant difference in amplitude between the groups studied; results do not corroborate this research. Sharma et al. (40) found similar results like the present study, comparing children aged between 10 and 19 years who practice sports and those who do not, using a Go/No-go characteristic paradigm. Using the Flanker Task, Ludyga et al. (48) evaluated the effect of practicing ten weeks of exercise training in children between 9 and 10 years old. The results demonstrated no difference between the pre-and post-intervention moments in the amplitude and latency variables. In another study, the same author employed the Go/No-Go paradigm with children between 9 and 13 years old with at least six months of Judo practice with non-practitioners and found amplitude results similar to those of the present study. However, the latency results differ due to the difference in practice and exercise time. Of the studies cited, none classified participants according to biological maturation, making direct comparison of results difficult.

Cognitive development across the lifespan is associated with changes in the neuroanatomical and neurochemical network (49–51). Some research using magnetic resonance imaging has shown that the Parietal Lobe matures earlier than other regions (52–55); there is ample evidence that the parietal lobe undergoes significant development from the age of 8 to 15, which directly implies the development of visual processing networks in children (56,57). Previous research has shown that the maturation of frontal and parietal networks is associated with maturational processes of cognitive control, which contributes to the development of working memory and attention allocation (58,59), and therefore, as age passes, there is an increase in activation of the parietal cortex related to the functional maturation of working memory and attention (60,61). This may explain why, the present study found a significant difference between PHV (Level −2 and Level −3) suggesting that Karate practice facilitates the maturation of this area. This finding justifies the observed interaction between Group and PHV.

### 4.4. Associative Somatosensory Cortex: Brodmann Area 5 (Cz)

Brodmann area 5, located in the superior parietal lobe immediately posterior to the primary somatosensory areas, is directly involved in visuospatial processing, including spatial perception. It also plays a crucial role in various types of working memory (motor, visual, auditory, emotional, and verbal), as well as visuospatial memory and visuomotor attention (62,63). This region is involved in several processes during the preparation and execution of motor action (64), containing a robust internal representation of cutaneous sensations from the fingers and movement (65,66). Lower amplitude and latency values suggest a specific capacity for change in the region’s dynamics that participates in motor elaboration, planning, and final action (67,68). Specifically, a smaller P300 amplitude suggests greater participants’ engagement in detecting target stimuli (69). Lower latency values may indicate enhanced efficiency in visuospatial processing, working memory, visuospatial memory, and visuomotor attention (70).

The results of the present study showed interaction in the P300 amplitude for the PHV factor. The lower amplitude values found in participants closer to the PHV (Level - 2) suggest that the practice of Karate could modify the dynamics of this region. Consequently, the lower P300 amplitude in these participants suggests greater engagement in detecting target stimuli. Hillman et al. (71), conducted research comparing children who exercise, children who do not exercise (9 to 11 years old), adults who exercise and adults who do not exercise (19 to 20 years old) using the Oddball paradigm, when comparing exercising and non-exercising children (9 to 11 years old), and adults (19 to 20 years old) using the Oddball paradigm, founding lower amplitude values and lower latency values in this region in children and adults who exercise, with the same dynamics found in previous studies by the same author. However, these studies were carried out with adults who practice sports and those who do not practice sports (72,73), which makes us believe that the practice of exercise and the more significant development of the area in question, leads to lower amplitude values and lower latency values, which in a way helps us understand the results of this study, as an interaction was found between the factors Group and PHV, with a significant difference for PHV (Level −2 and Level - 3). Even though no significant difference was found for Group, the experimental group presented slightly lower amplitude values than the non-Karate practitioners group. Using the Flanker Task, Lind et al. (47) found higher latency values and lower amplitude values in children who play football when compared to children who practice walking aged between 11 and 12 years, which somewhat corroborate our study. Ludyga et al. (74), used a Go/No-go characteristic paradigm to compare children who exercise with non-exercisers aged between 9 and 13 years and found lower amplitude and latency values in the exercise group, with results similar to those of the present research. The author explains his results by defending the idea that the Go/No-go task is mediated by a more favorable allocation of attentional resources and more effective conflict monitoring due to greater activation of this region in children who practice physical exercise, which did not occur in this study, even though a Go/No-go paradigm was also used, perhaps the shorter intervention time with the exercise was responsible for the difference a difference.

### 4.5. Secondary Motor Cortex: Brodmann Area 8 (Fz)

Located immediately anterior to the premotor cortex (Brodmann Area 6), it contains the Superior Fontal, Middle, and Medial Frontal Lobe Gyri (62,75,76). This region plays a key in several higher cognitive functions, including decision-making, planning, attention processes, and working memory (42,77). Furthermore, the primary activity of this region reflects the orchestration of thoughts and actions following internal goals (78). The findings of the present study indicated that there was no interaction between the factors. However, the results showed a main effect for the Group and PHV factors on P300 amplitude and latency.

The lowest P300 amplitude was found in the Karate practitioners group compared to the non-Karate practitioners group, and the participants were classified as PHV Level −3. This region is central to working memory and inhibitory control (79,80). According to Gordeev (81), the higher P300 amplitude value may be related to a greater state of alertness due to the continuous repetition of the task. And the complexity of the task may also significantly influence it. The central premise of the P300 pattern about amplitude variation is that this parameter is sensitive to attentional engagement; in other words, it will be greater or lesser depending on the amount of variation in attention dedicated to executing the task and new information resources (67,82), as evidenced in our results. We also observed a greater P300 latency in participants in the Karate practitioners group, and participants were classified as Level −3 of the PHV. In short, latency is the recognition of the stimulus classification speed resulting from discriminating one event from another concerning a specific context (62). The current P300 latency result indicates better information processing for executing the Karate practitioners group’s task as expected. These findings indicated greater neural efficiency in this group and greater engagement of attentional resources and working memory to perform the task. Li et al. (83) observed similar results using the Oddball paradigm in children between 8 and 12 years who underwent a 6-month physical exercise intervention, finding lower amplitude and latency values in the post-intervention. Sharma et al. (40) found similar results in the previous study comparing sports practitioners with non-practitioners. The age range studied was 10 to 19, and a Go/No-go sound stimulus characteristic paradigm was used. The research by Mora-Gonzalez et al. (84) and Raine et al. (85), when comparing exercisers and non-exercisers in children aged 8 to 11 years old and 6 to 11 years old, respectively, using a paradigm visuomotor, corroborate our results finding lower amplitude and lower latency in the group of exercisers.

Studies that compared exercisers and non-exercise children between 9 and 11 years old using visuomotor paradigms found higher amplitude values and lower latency values in physical exercisers. Such studies suggest that regular exercise can be associated with increased neuroelectric activation related to attention allocation and working memory (71,86). A factor that may influence study results is the maturation of this area, which, according to several studies. occurs later. These studies evaluated individuals every two years from ages 4 to 21 using magnetic resonance imaging (52–55) and found that specifically the prefrontal cortex exhibits non-linear development and involves neurogenesis and synaptogenesis processes (53). As none of the studies cited here investigated the neurodevelopmental stage of the area in question, it is possible that the samples from these studies are not homogeneous in this regard, directly interfering with the results. The present study was concerned with homogenizing the studied sample through PHV classifications. Possibly for this reason, the latency results found are not only different from other studies but also difficult to compare.

### 4.4. Limitations

This study was limited to examining only two biological maturation ranges according to PHV (Level −2 and Level −3); it would be interesting to expand the sample to include children from other maturation ranges. It would also be beneficial to study other sports, including various martial arts, to determine how different types of exercise or sporting practices influence attention and memory, and how they compare to one another. Another limitation is thecross sectional comparison research design, a longitudinal research would help to better understand the variables studied. We can also consider other types of exercise, mainly coordinative ones, which are better for developing cognition, according to acute data from Budde et al. (87) research and supported by longitudinal data Koutsandreou et al. (88). Another limitation is the condition with no karate. We should in a next step compare our results with exercise non inferiority trail (89).

## 5. Conclusion

The hypotheses of lower latency and amplitude values in the Karate practitioners group was partially confirmed. In general, it is possible to conclude that participants in the Karate practitioners group exhibited electrocortical measures corresponding to greater efficiency in decision-making and attention processes, motor planning, working memory, attention allocation, motor execution, and greater attentional engagement. It was also demonstrated that, despite the children being at very close chronological ages, their biological maturation differed. This difference interfered with some of the results of the variables studied since the children classified at Level −2 of the PHV had lower amplitude and latency results than those classified at Level −3. Therefore, biological maturation is an important variable to be controlled in future studies. However, further studies are required to expand this knowledge.

